# A replication hijacking mechanism for Tn*3*-family replicative transposition

**DOI:** 10.64898/2026.05.30.728999

**Authors:** Emilien Nicolas, Cédric A. Oger, Claire Stulemeijer, Nicolas Aryanpour, Ophélie d’Udekem d’Acoz, Nathan Nguyen, Michael Chandler, Bernard F. Hallet

## Abstract

Transposition of all classes of transposable elements generates DNA intermediates that must be processed by the host to be effective. However, the mechanisms whereby transposons communicate with cellular DNA-processing machineries remain poorly investigated. Here, we provide convergent genetic and biochemical evidence that replicative transposition of the Tn*3*-family transposon Tn*4430* is strictly coupled to replication of the target. Blocking target replication abolishes transposition, while blocking replication of the transposon donor molecule has no effect. Furthermore, the insertion preference of Tn*4430* was found to be altered by the direction of replication and potential replication impediments within the target, suggesting a functional link between the integration mechanism and replication fork progression. *In vitro*, the transposase TnpA was found to specifically bind to fork-like DNA structures that mimic replication intermediates. Compared to linear DNA fragments, these structures are efficient substrates for TnpA-catalysed end joining. Strand transfer occurred immediately downstream of the fork, poising the transposon for replication. Together, the data suggest a mechanism in which the transposon targets DNA replication intermediates to directly recruit the host replication machinery at the time of transposition. This “replication hijacking” mechanism contrasts with classical “replication hiring” mechanisms during which replication is recruited after strand transfer.

**SIGNIFICANCE STATEMENT:** Bacterial transposons of the Tn*3* family constitute a threat for human health, being continuously involved in the emergence and spread of new antimicrobial resistances amongst pathogens. The success of these elements is based on their replicative mode of transposition allowing them to produce a new copy of themselves whenever they move. Here, we provide evidence that, rather than recruiting the host replication machinery after integration as is proposed in textbook models for replicative transposition, Tn*3*-family transposons directly transpose into ongoing replication intermediates. We propose that this “replication hijacking” mechanism provides a means of synchronizing transposition with cell replication activity, thus optimizing the movement of transposons with the expansion of bacterial populations.

**GRAPHICAL ABSTRACT:** 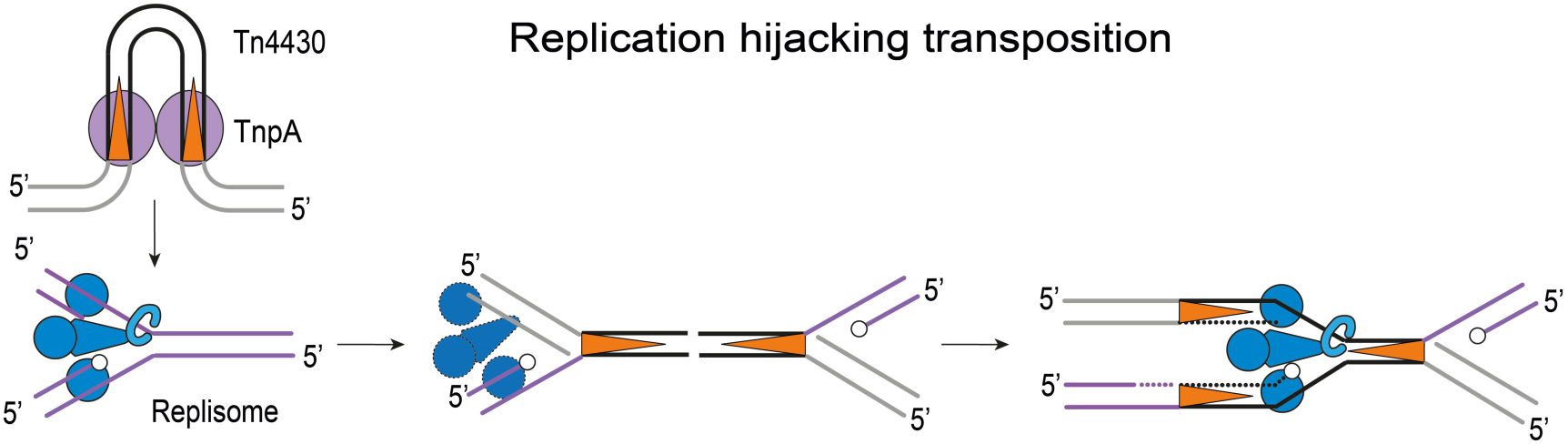

## INTRODUCTION

Transposons group a diverse range of mobile genetic elements that can propagate from one location to another within the genome of their host (1). The movement of these elements is mediated by a variety of mechanisms, some of which are conserved from bacteria to higher eukaryotes, evidencing the importance of transposable elements in the shaping and evolution of genomes. In prokaryotes, transposons are the driving force of the mobilome, allowing bacterial communities to face environmental challenges by re-assorting useful phenotypic traits through horizontal gene transfers (2,3). In eukaryotes, transposable elements play an architectural role in the structural and functional organization of chromosomes, being intimately implicated in the creation of complex regulatory networks and other genetic innovations (4,5).

This evolutionary success necessarily requires mutual adaptations between transposable elements and their hosts (6). On the one hand, transposition must be tightly regulated to preserve the host genome from deleterious DNA rearrangements. On the other hand, transposable elements rely on the host cellular machineries to process DNA intermediates that are generated by the transposition reaction. For transposons that move by a simple “cut-and-paste” mechanism, completion of transposition merely requires host-mediated repair of small gaps resulting from staggered DNA cuts that are catalyzed by the transposase upon insertion into the target. Closing the gaps typically generates short target site duplications (TSD) that flank the transposon after integration (7,8). However, different classes of transposons use more elaborate replicative mechanisms that duplicate the entire element as an integral part of the transposition process. This can occur by reverse-transcribing a primary transcript into a cDNA copy (class I retrotransposons), or by recruiting the cell replication machinery at a specific stage of direct transposition between a donor and a target DNA molecule (replicative class II DNA transposons) (5,7–11). In either case, the mechanisms whereby the transposition machinery and endogenous DNA-processing enzymes coordinate their activities remain poorly explored.

Here, we address this issue taking the transposon Tn*4430* from *Bacillus thuringiensis* as a model system for studying the replicative transposition mechanism of the Tn*3* family (12,13). Members of this family are widespread and function as mobile platforms for a broad range of passenger genes, including a number of antibiotic resistance genes (12,14,15). They are notably involved in the recent outbreak of carbapenem-resistant enterobacteriacae and the even more recent dispersal of resistance to colistin, which seriously compromises the use of these two antibiotics as a last-resort defense against multi-resistant infections (16–19).

The effectiveness of Tn*3*-family transposons largely relies on their “paste-and-copy” or “copy-in” mechanism of replicative transposition during which a new copy of the transposon is generated upon integration into a new target (12). This mechanism has been studied in detail for bacteriophage Mu which uses replicative transposition to multiply its genome during lytic growth (11,20,21). At the onset of Mu transposition, DNA cleavage and joining reactions catalyzed by the transposase MuA generates a branched strand transfer product with Y-shaped structures resembling a replication fork at both ends of the phage genome. DNA pol III then initiates replication at one of these forks following a cascade of events leading to the disassembly of the transposition complex (or transpososome) and its replacement by the host replisome (22–24). By re-routing cellular replication to its profit, this “replication-hiring” mechanism ensures efficient replication of Mu, allowing to produce up to 100 copies of its genome per infected cell (11,21).

Our data suggest that transposons of the Tn*3* family employ a conceptually distinct “replication-hijacking” mechanism during which the transposition complex directly recruits the replication machinery by mediating transposition into ongoing replication intermediates. This alternative mechanism is supported by genetic evidence demonstrating that contrary to Mu, target replication is an absolute requirement for Tn*4430* transposition and that it influences its integration preference *in vivo*. In addition, the Tn*4430* transposase, TnpA, was found to specifically bind to replication fork-like DNA molecules *in vitro* and to use them as a substrate for the strand transfer reaction that joins the transposon ends to the target. We propose that this mechanism allows synchronization of transposition with DNA replication, thus promoting the movement of transposons in bacterial populations that are active and expanding.

## MATERIAL AND METHODS

### Bacterial strains, Plasmids and oligonucleotides

Strains, plasmids and oligonucleotides used in this study are listed in Table S1

### Transposition into temperature-sensitive replication targets

The mobile cassette Mini-TnKm consists in a kanamycin resistance gene flanked by the 38-bp terminal inverted repeats (IRs) of Tn*4430*. The integration pattern of Mini-TnKm into the temperature sensitive replication plasmids pGB2T°S, pGINC5LacO, pGIENAsp and pGINC5lacO-α was determined by plasmid rescue assay as previously described (25). Briefly, LacI-free HB101Δ*lacI E. coli* cells harboring the mini-TnKm donor plasmid pGIML027, the TnpA expression vector pGIEN001.1, and the target were grown at replication-permissive T° (30°C) and cointegrate products were selected by spreading the cultures on selective LB plates at non-permissive T° (42°C). Between 24 and 30 cointegrates were recovered for each target and resolved by restriction (using *Ecor*V and *Bsa*AI) and ligation to remove one of the two copies of Mini-TnKm. The position and orientation of the remaining copy within the target was determined by DNA sequencing using the outward-facing ENminiTn01 and ENminiTn03 primers internal to Mini-TnKm (Table S1). The 5-bp TSDs of insertion sites were aligned along with 10 bp of flanking DNA according the orientation of Mini-TnKm within the target. Target sequences that were recovered multiple times were included only once in the alignment. For each target, a sequence consensus was generated using Weblogo (http://weblogo.berkeley.edu).

### Transposition with conditionally replicating target and donor plasmids

To determine the role of target replication in Tn*4430* transposition, electro-competent cells of the λ phage lysogenic strain CSH50λCI harboring the TnpA (or TnpA^D751N^) expression vector pGIEN2.1 (or PGIEN2.1Mtc) and the Mini-TnKm donor plasmid pGIML027 were transformed with the λ-derived target plasmid PCB104. In the reciprocal experiment to assess the role of donor replication, the λ-derived plasmid pGIML028 was used as a donor and pBG01 was used as a target. Since CSH50λCI cells express the λ CI repressor, the transformed λ-derived target or donor plasmid cannot initiate replication, and the only way to rescue the chloramphenicol resistance is by forming a transposition cointegrate. Electroporation mixes were incubated with 0,02% L-arabinose for 2h at 37°C to express TnpA and transposition products were selected by plating the cells on Ampicillin (Ap), kanamycin (Km) and chloramphenicol (Cm) for experiments performed with the pCB104 target and the pGIML027 donor; or Ap, Km and tetracycline (Tc) for experiments performed with the pGIML028 donor and the pBG01 target (Fig. 2A). Transposition frequencies were measured by dividing the number of Ap^R^/Km^R^/Cm^R^ (or Ap^R^/Km^R^/Tc^R^) clones by the total number of Ap^R^/Km^R^ (or Ap^R^/Tc^R^) cells in the electroporated samples, respectively.

To validate the transposition partners under conditions where both the target and donor molecules can replicate, CSH50 cells containing either one or the other combination of the three plasmids were plated on selective medium supplemented with 0.02% L-arbinose. Plasmid DNA was then extracted from a pool of ∼200 colonies, and cointegrates formed with the λ-derived pCB104 target or pGIML028 donor were selected by transforming the CI-expressing strain CSH50λCI. In this case, transposition frequencies were expressed as the number of Km^R^+Cm^R^/Km^R^ transformants obtained with the pGIML027 donor and PCB104 target, and the number of Km^R^+Tc^R^/Tc^R^ clones obtained with pGIML028 donor and pBG01 target, respectively (Fig. 2A).

### Proteins and DNA substrates

TnpA proteins were purified by nickel affinity chromatography using a C-terminal (His)_6_ tag as previously described (13,26). Purified fractions were dialyzed against TnpA storage buffer (50 mM Tris-HCl pH 8.0, 1M NaCl, 30% glycerol) typically giving a final TnpA concentration of ∼0.3 – 1M Fluorescently-labeled target DNA substrates were assembled by annealing complementary oligonucleotides as specified in tables S1 and S2. 5’-Cyanidin 3- (Cy3-), 5’- Cyanidin 5- (Cy5-) and 5’-HiLyte 647 (H647-) labeled oligonucleotides were puchazed from Eurogentec. 5’-^32^P labeling of oligonucleotides was performed with [γ^32^P]-ATP (Perkin-Elmer) and T4 polynucleotide kinase (New England Biolabs). 70-bp pre-cleaved donor substrates containing the Tn*4430* IR sequence were assembled by annealing 5’H647-EN100-3’-1 with EN100-5’ (Table S1 and S2). 120-bp pre-cleaved donors were PCR-amplified from the Mini- TnKm-containing pGIML027 plasmid using 5’-^32^P-COD2 or 5’-Cy3-END2 and END1 primers (table S2), followed by N*co*I cleavage to mimic IR end processing by TnpA (13). Strand transfer products (STP) used as a starting substrate in the disintegration reaction were pre-assembled by annealing three oligonucleotides (Table S1 and S2). DNA substrates were separated on 8%

(vol/vol) polyacrylamide gels in TBE buffer (88 mM Tris base, 88 mM boric acid, and 2 mM EDTA) and eluted from a gel slice in TES buffer (10 mM Tris HCl pH 8.0, 1 mM EDTA, 300 mM NaCl) prior to be ethanol-precipitated and resuspended in TE buffer (10 mM Tris HCl pH 8.0, 1 mM EDTA).

### Electrophoretic mobility shift assays (EMSA)

EMSA reactions (20 μL) were performed by mixing TnpA (75–300nM) with ∼800ng of DNA substrate in standard binding buffer [50mM Tris HCl pH 8.0, 1 mM DTT, 5 mM EDTA, 50 mM NaCl, 10 mg/mL BSA, 10% (vol/vol) DMSO]. Reactions were incubated at 34 °C for 15 min and run on 4% (vol/vol) polyacrylamide gels in TBE buffer at 4 °C. Gels were imaged with a Typhoon X biomolecular imager (Amersham) and quantified with ImageJ.

#### Integration and disintegration *in vitro*

Integration of Tn4430 IR ends into synthetic target molecules was performed by incubating TnpA (150-300 nM) with 400ng of H647-labeled or ∼100 cps (count per second)/µL of P32- labeled pre-cleaved IR ends and 200-800ng of fluorescently labeled target in binding buffer supplemented with 10% (vol/vol) PEG 6000 and 5 mM MnCl_2_ or MgCl_2_ instead of EDTA. Disintegration was analyzed using pre-assembled Cy5/Cy3-labeled STP as the initial substrate (Table S2). Reactions were incubated at 34°C for 1 to 16 h and deproteinized with proteinase K (600 μg/ml) (Sigma) in 0.2% SDS, 12.5 mM EDTA (pH 8.0) followed by phenol-chloroform extraction and ethanol precipitation or directly purified with Monarch kit (New England Biolabs). Products were resuspended in TE buffer [10 mM Tris HCl pH 8.0, 1 mM EDTA] and analyzed by electrophoresis on non-denaturing 7.5% polyacrylamide gels in TBE buffer, or denatured by heat treatment (95°C, 10 min) in formamide loading buffer [80% (vol/vol) formamide, 1 mM EDTA pH 8.0, 10 mM NaOH, bromophenol blue] and separated on denaturing 8% polyacrylamide, 25% (vol/vol) formamide gels in TBE buffer. Fresh gels were scanned with an Ettan DIGE (GE-Healthcare) or Typhoon X (Amersham) imagers for fluorescence scanning and dried and autoradiographed with a Pharos FX scanner (Bio-Rad) for ^32^P detection.

## RESULTS

### Regional versus sequence-specific integration of Tn*4430*: the replication targeting hypothesis

In a previous study, we showed that the presence of multiple copies (3 or 5) of the *Escherichia coli lacO* operator downstream of the replication origin of a pSC101-derived plasmid strongly altered the insertion preference of Tn*4430* as determined by PCR assay (25). To obtain further insight into the influence of the *lacO* repeat on Tn*4430* targeting, independent insertions of a Tn*4430*-derived mobile cassette (Mini-TnKm) into the *lacO*-free and 5-*lacO*-containing plasmids were mapped by DNA sequencing (Fig. 1). In the *lacO*-free plasmid, 19 of the 24 sequenced insertions were clustered in the intergenic region (hotspot 1, HS1) between the replication operon *rep* and the Sp/Sm resistance gene of the plasmid (Fig. 1A). In the target carrying the 5-*lacO* repeat, this regional preference was shifted to a distinct hotspot region (HS2) spanning the region from the unidirectional replication plasmid origin to the 5-*lacO* array, representing 18 of 29 insertions over 696 bps (Fig. 1B). The insertion pattern obtained for an equivalent target in which the 205-bp 5-*lacO* array has been replaced by a non-specific (NS) DNA fragment from the TEM1 β-lactamase gene of pBR322 (Table S1) was similar to that obtained for the *lacO*-free plasmid, with suppression of HS2 and a higher density of insertions around HS1 (Fig. 1C)

**Figure 1.**
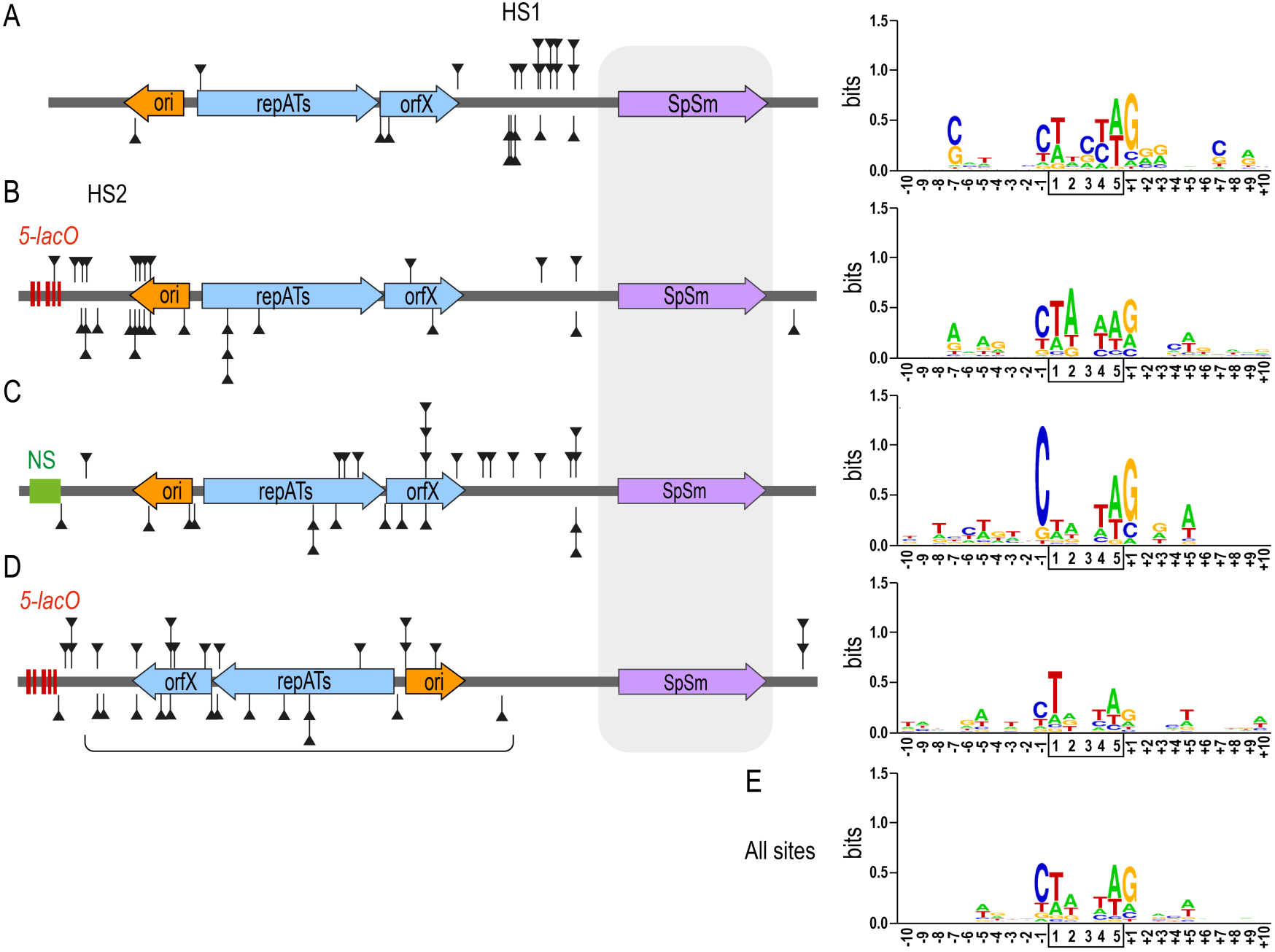
Differential DNA targeting by Tn*4430*. Independent Mini-TnKm insertions were mapped into the pSC101-derived thermosensitive plasmid pGB2Ts (A), pGINC5lacO containing an array of 5 copies of the *lacO* operator (red bars) (B), pGIENAsp in which the 5-*lacO* repeat has been substituted for a 250-bp non-specific fragment (NS, green box) (C), and pGINC5lacO-α in which a 2.3-kb fragment (bracket) carrying the replication operon (rep, blue arrows) and the unidirectional replication origin of the plasmid (ori, orange arrow) has been reversed with respect to the 5*-lacO* repeat (D). Mini-TnKm insertions for which transcription of the Km gene is co-directional with replication are indicated by black triangles below the plasmid maps, while triangle above the maps show insertions with the opposite orientation. Regions corresponding to hotspot 1 (HS1) and 2 (HS2) are indicated. The region containing the SpSm gene (purple arrow) is refractory to insertions because of selection of transposition cointegrates on spectinomycin (shaded aera). Logo plots of the target consensus obtained for each plasmid are shown to the right, and the generic consensus obtained for all insertions is shown at the bottom (E). positions of the 5-bp TSDs are boxed, and positions of the flanking sequences are numbered ±1 to ±10.

Since these experiments were performed in a Δ*lacI E. coli* host, formation of HS2 in the 5-*lac0*-containing target is independent of the binding of the LacI repressor to *lacO*. In addition, alignment of the insertion sites revealed a weak signature with the consensus 5’- CTnnnAG-3’, whatever the target (Fig. 1.). We therefore conclude that Tn*4430* target site preference is primarily dictated by structural and/or functional DNA features rather than by primary sequence recognition, and that the specific presence the *lacO* repeat in the 5-*lacO* plasmid exerts a polar effect on the adjacent HS2 region to make it more attractive for the transposon. In particular, the 20-bp *lacO* sequence has a pseudo-palindromic organization that can potentially form secondary DNA structures, such as hairpins or cruciforms, which are known to affect replication (27–29). The repetition of such structures immediately downstream of the unidirectional replication origin of the plasmid might thus generate a specific zone in which DNA structures associated with replication transiently accumulate. We hypothesize that if these replication intermediates provide a positive signal for Tn*4430* integration (*e.g*., by interacting with the transposition machinery), their accumulation in HS2 could account for the observed preference. According to this hypothesis, transposition into secondary hotspots like HS1 may arise from other processes interfering with DNA replication, such as collisions between replication and transcription from the *rep* operon (30)(Fig. 1).

Further support for the idea that target replication affects Tn*4430* integration specificity comes from the observation that transposition into a plasmid where the replication origin has been reversed and placed distal to the 5*-lacO* repeat gave yet another pattern with broadly distributed insertions throughout permissive target regions of the plasmid (Fig. 1D).

### Replication of the target is essential for transposition

The possibility that replication intermediates are preferred Tn*4430* targets implies that target DNA must undergo active replication for efficient transposition to occur. To test this idea, we set up a plasmid rescue assay with conditionally replicating target and donor molecules (Fig. 2). In this assay, a λ-lysogenic *E. coli* strain CHS50λ harboring a TnpA expression vector and a compatible Mini-TnKm donor plasmid was electro-transformed with a target molecule that functions with the replication origin of bacteriophage λ (Fig. 2A). Since CHS50λ cells express the λ CI repressor, replication initiation of the transformed λ-derived target is inhibited unless it is rescued by forming a transposition cointegrate with the donor plasmid (Fig. 2A, left). In a reciprocal experiment, TnpA-expressing CSH50λ cells containing the target plasmid were electro-transformed with a λ-derived plasmid which, this time, contained mini-TnKm and thus acted as a donor in transposition (Fig. 2A, right).

**Figure 2.**
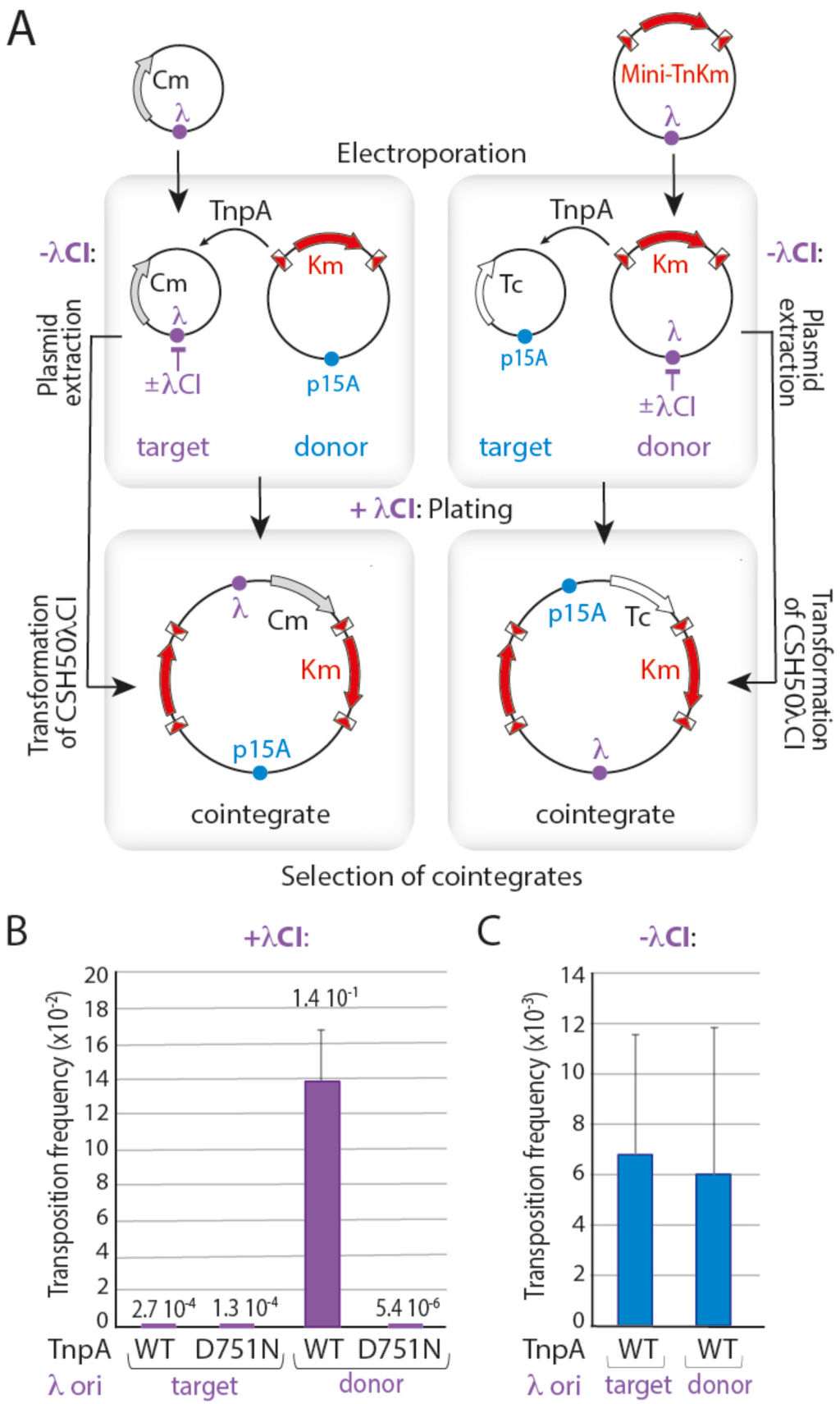
Role of target and donor replication during transposition. **(A)** Suicide plasmid transposition assay. Cells of the λ lysogenic E. coli strain CSH50λCI harboring a p*Ar*a-driven TnpA^WT^ or TnpA^D751N^ expression vector (not shown) and either a p15A-derived donor (left) or target plasmid (right) were respectively electro-transformed with a partner target (left) or donor plasmid (right) containing the λ phage origin of replication (λ). As the cells express the λ CI repressor (+λCI), replication of the electroporated plasmid is inhibited but can be rescued by transposition of Mini-TnKm. Cointegrate products were recovered by selecting clones on kanamycin (Km) and chloramphenicol (Cm) for the suicide target (left), or on Km and tetracycline (Tc) for the suicide donor (right). **(B)** Transposition frequencies obtained with the λ-derived target or donor suicide plasmid are calculated by normalizing the number of Km^R^/Cm^R^ or Km^R^/Tc^R^ clones to the transformation rate of the competent cells, respectively. **(C)** Transposition under replication-permissive conditions of λ-derived plasmids (-λCI). The TnpA expressing vector and the two pairs of target/donor partners were grown in the non-lysogenic strain CSH50. Plasmid DNA was then extracted and used to transform CSH50λCI to select for cointegrates. Transposition frequencies are expressed as the ratio of Km^R^/Cm^R^ or Km^R^/Tc^R^ clones over the number of Km^R^ or Tc^R^ transformants obtained for the p15A-derived partner, respectively.

With the non-replicating donor, the frequency of co-integrate formation by wild-type TnpA (TnpA^WT^) (1.4 10^-1^) was 4 to 5 orders of magnitude higher than the background measured for the catalytically inactive mutant TnpA^D751N^ (5.4 10^-6^) (13) (Fig. 2B); whereas with the non-replicating target, transposition activity was essentially undetectable, the frequency of Km^R^/Cm^R^ colonies measured for TnpA^WT^ (2.7 10^-4^) being in the same range as for TnpA^D751N^ (1.3 10^-4^)(Fig. 2B).

To verify that the results obtained with the donor and target suicide plasmids were indeed the result of their ability to replicate or not, transposition was examined under conditions where both partners were capable of autonomous replication (Fig. 2C). To this end, the plasmids used in either version of the suicide experiment were grown in the non-lysogenic strain CSH50 that is permissive for λ replication. Plasmid DNA was then extracted from the cells and used to back-transform the non-permissive lysogenic strain CSH50λ in order to select for cointegrate products (Fig. 2A). In this experiment, the transposition frequency was in the same range, whether the λ-derived replicon was the target or the donor plasmid (Fig. 2C). This demonstrates that both pairs of plasmids are efficient substrates for TnpA-mediated transposition, and that the absence of transposition observed with the λ-derived target resulted from replication inhibition. Replication of target DNA is thus essential for Tn*4430* transposition while replication of the donor is not, demonstrating a direct role of target DNA replication in the integration mechanism.

### The Tn*4430* transposase specifically binds to replication fork-like structures *in vitro*

The regional preference of Tn*4430* and its dependence on target DNA replication raises the possibility that the transposase TnpA interacts with replication intermediates to mediate integration. To test this hypothesis, electrophoretic mobility-shift assay (EMSA) was used to examine the ability of TnpA to bind potential target DNA molecules with different structures *in vitro* (Fig. 3). This analysis was performed with TnpA^WT^ and two deregulated mutants, TnpA^S911R^ and TnpA^W24R/A174V/E740G^ (hereinafter referred to as TnpA^3X^), which were initially identified for their impairment in target immunity, a yet ill-defined regulatory mechanism whereby Tn*3*-family transposons avoid inserting multiple times into the same target (25,31). TnpA^S911R^ and TnpA^3X^ displayed promiscuous target selectivity *in vivo*, mediating transposition into an immune target at a higher frequency than TnpA^WT^ and proved to be hyperactive in different biochemical assays *in vitro* (13,15,31). At the structural level, hyperactivity of both TnpA mutants correlates with their ability to form an activated paired-ends complex (PEC) consisting of a TnpA dimer bound to two terminal inverted repeats (IR), which TnpA^WT^ cannot form spontaneously (13,15,32).

**Figure 3.**
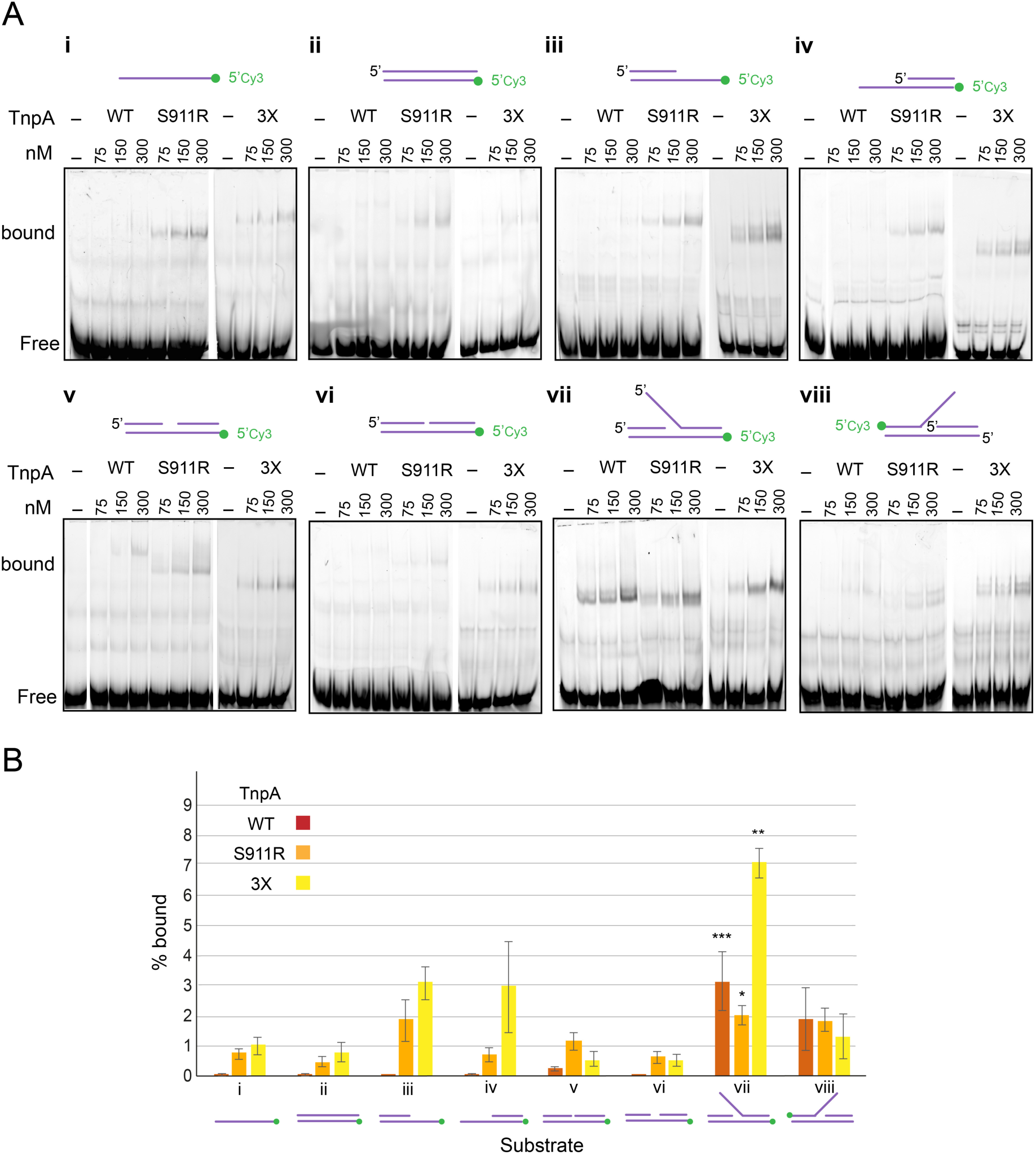
TnpA binding to different target structures *in vitro*. **(A)** 5’-Cyanin 3- (Cy3-) labelled DNA substrates corresponding to ssDNA (i), different forms of dsDNA duplexes (ii to vi) or branched fork- like DNA molecules (vii and viii) were assembled using the oligonucleotides listed in Supplementary Table S2, and incubated with increasing amounts of TnpA^WT^ (WT), TnpA^S911R^ (S911R) or TnpA^3X^ (3X) as indicated. Binding reactions were run on a 4% polyacrylamide gel and analyzed by fluorescence scanning. **(B)** For each substrate, the percentage of bound DNA was measured at the highest TnpA concentration (300 nM). Error bars indicate standard errors obtained from 3 to 4 independent measurements per substrate. Binding preference was established by anova analysis with p ≤ 0.05 (*), ≤ 0.01 (**) and ≤ 0.001(***) (see Data availability).

TnpA^WT^ binding to single-stranded DNA or to different DNA duplexes bearing blunt or recessive ends, a 5-nt gap or a single-strand nick was not or barely detectable, confirming its low non-specific DNA binding activity (Fig. 3, substrates i to vi) (13,32). In contrast, TnpA^WT^ binding was significantly increased on a branched DNA molecule having the specific structure of a replication fork with a 3’OH-ended leading strand and a 5’-single-stranded flap corresponding to the lagging strand template (Fig. 3, substrate vii). To a lesser extent, TnpA^WT^ also binds to a DNA fork with the opposite polarity (*i.e*., with 5’-end instead of a 3’-end at the branch point), a structure that is incidentally found at arrested replication forks *in vivo* (33,34)(Fig. 3, substrate viii). Presumably reflecting their deregulated phenotype, TnpA^S911R^ and TnpA^3X^ were found to be more promiscuous than TnpA^WT^, giving detectable binding with all DNA substrates tested, but still with a marked preference for DNA forks compared to linear duplex (Fig. 3). Both mutant TnpAs also displayed enhanced binding to the linear fragment harboring a recessed 3’OH end as in typical DNA synthesis intermediates (substrate iii) and to a lesser extent, to the 5’-recessed fragments as found in the reversed fork (substrate iv). Thus, despite low or no affinity for nonspecific DNA sequences, TnpA specifically interacts with DNA structures that mimic DNA synthesis intermediates, suggesting that these structures may play a role in DNA targeting by Tn*4430*.

### Fork-like DNA structures are preferred substrates for TnpA-catalyzed strand transfer *in vitro*

To determine whether particular DNA structures are targets for TnpA-catalyzed strand transfer, the above Cy3-labelled DNA substrates were mixed with a 5’Cy5-labelled pre-cleaved donor IR end and incubated with purified TnpA^WT^ or hyperactive TnpA^S911R^ *in vitro* (Fig. 4). In the pre-cleaved donor, the external 3’OH group of the IR is exposed to mimic TnpA-catalyzed DNA cleavage, which has been shown to increase the rate of strand transfer by promoting active PEC formation (13). The reactions were deproteined and analyzed by electrophoresis in a native 7.5% polyacrylamide gel followed by fluorescence scanning (Fig. 4).

**Figure 4.**
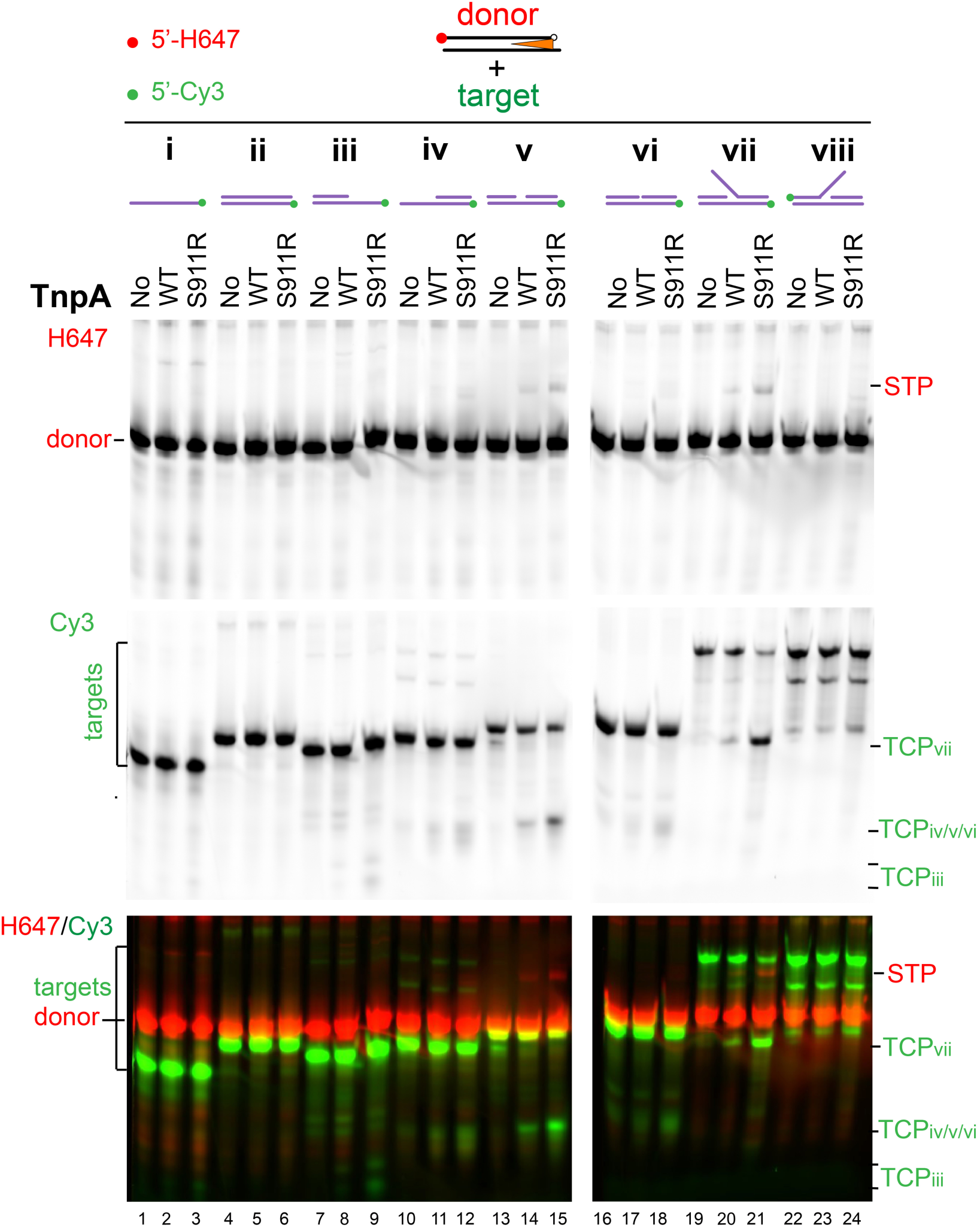
Integration of Tn4430 IR end into replication-derived DNA structures *in vitro*. A 5’-Hylite 647- (H647-) labeled pre-cleaved donor IR (orange triangle) was incubated with Cy3-labeled target substrates I to vii (see also Fig. 3) in the presence of TnpA^WT^ or TnpA^S911R^ as indicated (No, no TnpA). The reactions were run on a 7.5 % polyacrylamide gel and scanned for H647 (upper panel) or Cy3 (middle panel) florescence, or both (lower panel). Positions of the H647-labeled strand transfer products (STP) and Cy3-labeled target cleavage products (TCP) are indicated (corresponding substrates in subscript).

Incubation of TnpA^WT^ and TnpA^S911R^ with the replication fork-like substrate (substrate vii), and to a lesser extent with the linear substrate containing a 5-nt gap (substrate v), gave a Cy5-labelled strand transfer product (STP) of increased size compared to the donor, consistent with the formation of a covalent bond between the pre-cleaved IR end and the target. STP formation coincides with the presence of a Cy3-labeled band migrating ahead of the target band, indicating the release of a target cleavage product (TCP). With both substrates, TnpA^S911R^ produced higher levels of STP and TCP than TnpA^WT^, which again is consistent with its higher propensity at forming the active complex (13,15,32). No integration into the fork was observed in the absence of cofactor or with the catalytic mutant TnpA^S911R/D751N^. The reaction did not occur when Mn^2+^ was replaced by Mg^2+^, confirming that Mn^2+^ stimulates TnpA activity *in vitro* as is often the case for DDE/D polynucleotidyl transferases (Fig. S2) (13,35).

Time course analysis confirmed that the fork-like substrate (vii) is a more efficient target for strand transfer than the gapped substrate (v) giving faster and much stronger accumulation of TCP over time with both TnpA^WT^ and TnpA^S991R^ (Fig. S2). However, unlike TCP, STP accumulation was not linear and reached a plateau after ∼4 h of reaction, suggesting further processing by TnpA (see below). No or very weak STP or TCP bands were observed with the other substrates analyzed (i, ii, iii, iv, vi), including the reversed fork (viii), indicating that structures that better match bona fide replication intermediates provide better targets for strand transfer (Fig. 4).

### TnpA-mediated strand transfer into DNA forks takes place at a specific position, immediately downstream of the nascent 3’OH end

Since it is the best target for TnpA-mediated integration *in vitro*, strand transfer into the three-arm DNA fork (substrate vii) was characterized into further details using a P^32^-labeled donor IR end and a doubly labeled fork carrying a Cy5 and a Cy3 fluorophore at the 5’ end of the lagging and leading strand templates, respectively (Fig. 5A). As expected, incubation of the substrates with TnpA^WT^, TnpA^3X^ and TnpA^S911R^ generated a radiolabeled STP migrating above the donor fragment that was not observed with the catalytic mutant TnpA^S911R-D751N^ (Fig. 5B, lanes 2, 3 and 4). Formation of STP coincided with the release of a lower-molecular weight TCP retaining both fluorophores from the target (Fig. 5B, lanes 2’, 3’ and 4’). Again, the product bands were more intense for the deregulated TnpA^3X^ and Tnp^AS911R^ mutants than for TnpA^WT^. No STP nor TCP was detected when the donor or target substrates were incubated alone with TnpA, demonstrating that they arose from the transfer of the radiolabeled end into the fluorescent target and not from integration of the donor into itself or from simple cleavage of the fork in the absence of the donor end (Fig. S3).

**Figure 5.**
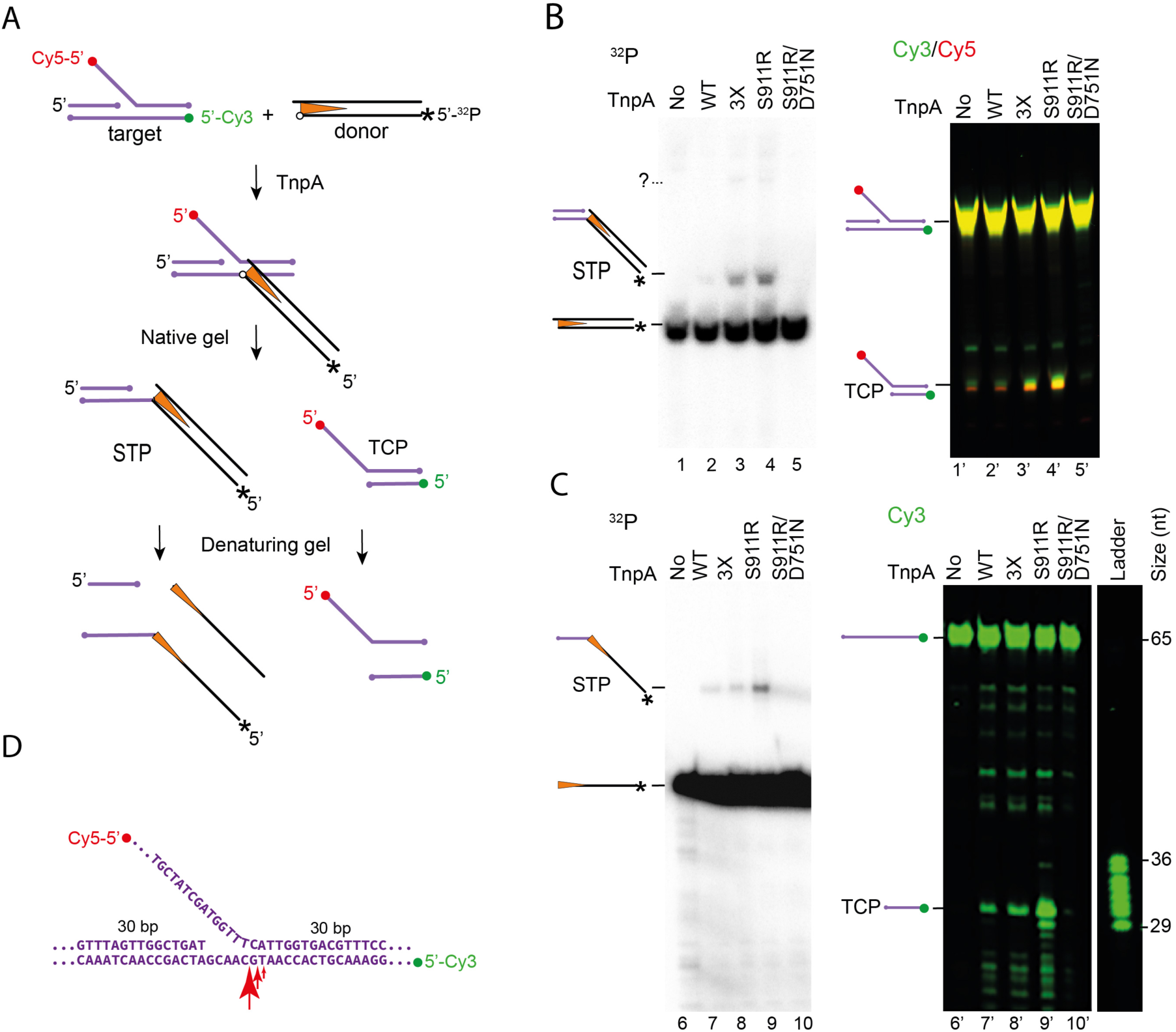
Specific integration into a DNA fork. (A) Integration of a 5’ ^32^P-labeled pre-cleaved IR end (*) was performed into a fluorescently labeled DNA fork containing a Cy5 label at the 5’ end of the lagging strand template (red dot) and a Cy3 label at the 5’ end of the leading strand template (green dot). (B) Reactions with the indicated TnpA variants were run on a native 7.5 % polyacrylamide gel and analyzed by phosphorimaging (left) to detect ^32^P-radiolabeled strand transfer products (STP), and by Cy5/Cy3 fluorescence scanning (right) to identify target cleavage products (TCP). “?” shows the position of a minor product observed with TnpA^3X^ and TnpA^S911R^ that could not be characterized further. (C) To map the integration site, reactions were analyzed by electrophoresis on an 8% polyacrylamide-25% formamide denaturing gel alongside a ladder of 5’Cy3-labeled oligonucleotides overlapping the branchpoint of the fork. Positions of radiolabeled and Cy3-labeled DNA strands resulting from denaturation of STP and TCP, respectively, are indicated. (D) Mapping of integration sites on the fork sequence with the larger red arrow showing the major site at the branch point. No, no TnpA.

To identify the DNA strand(s) targeted by strand transfer, integration was performed with forks that were 5’Cy3-labeled on each strand separately, and the products were analyzed on denaturing 8% polyacrylamide gels (Fig. S4). Fluorescence scanning of the gels revealed that only the leading strand template of the fork was cut by the reaction, releasing a specific fragment carrying the Cy3 fluorophore (Fig. S4). Mapping of the cleavage site on the doubly Cy5/Cy3-labeled fork showed that strand transfer occurred at specific positions at, or adjacent to, the branch-point of the fork (Fig. 5C). Interestingly, a similar pattern was observed when the branch point of the fork was shifted 5 bp downstream of its initial position, showing that the specificity of the reaction is primarily conferred by structural features of the target, rather than by a specific sequence (Fig. S5)

Thus, the results demonstrate that TnpA is capable of joining a Tn*4430* IR end to a specific position of a DNA fork, immediately downstream of the leading strand 3’OH end. No strand transfer occurred elsewhere in the fork. In particular, no obvious product band was observed for the leading strand or the lagging strand template as would be expected for concerted joining of two transposon ends into the target (Fig. 5 and Fig. S4). Faint radiolabeled bands, which likely resulted from rare integration at ectopic sites of the fork, were observed with TnpA^3X^ and TnpA^S911R^ (labeled “?” in Fig. 5B and Fig. S3), but could not be further characterized because they gave no detectable signal after denaturing gel electrophoresis.

### Processive integration and disintegration *in vitro*

As described above, prolonged integration reactions into the fork led to efficient cleavage of the target, while the corresponding STP appeared to accumulate to a lower extent (Fig. 4). This was confirmed by time-course analysis of integration, showing transient formation of STP and continuous accumulation of TCP over time (Fig. S2). To clarify the fate of the two partners of strand transfer, a 5’ Cy3-labelled donor IR was incubated with a fork that was this time Cy5- labeled at the 5’ end of the leading strand, and P^32^-labeled at the 5’ end of the template (Fig. 6A). Time course analysis performed with TnpA^S911R^ confirmed that the fork was gradually converted into a faster-migrating P^32^-labeled TCP as expected for strand transfer occurring at the branch point of the fork (Fig. 6B). Complete cleavage of the target was observed after 16h of incubation with TnpA^WT^ and both hyperactive mutants (Fig. 6C left). No cleavage was detected with TnpA^S911R-D751N^ or in the absence of the donor IR, confirming that cleavage of the target resulted from TnpA-catalysed strand transfer. However, fluorescence scanning of the gels revealed that the Cy3-labeled IR remained stationary with no apparent STP accumulation, while fork processing was correlated with the production of an additional Cy5- labeled fragment coming from the target (Fig. 6A and B). The most likely interpretation of this result is that primary STPs are processed further by TnpA, leading to disintegration as was previously reported for several transposases and polynucleotide transferases (36–41)(Fig. 6A). However, in this case, donor disintegration is likely not the true reversal of the integration reaction since donor excision does not reseal the initial target, but converts it to the observed disintegration product (DP). Supporting this scenario, incubation of preassembled STP led to excised IR and DP accumulation with all three TnpAs, yet with a faster kinetics for the two deregulated mutants TnpA^S911R^ and TnpA^3X^ (Fig. 6C).

**Figure 6.**
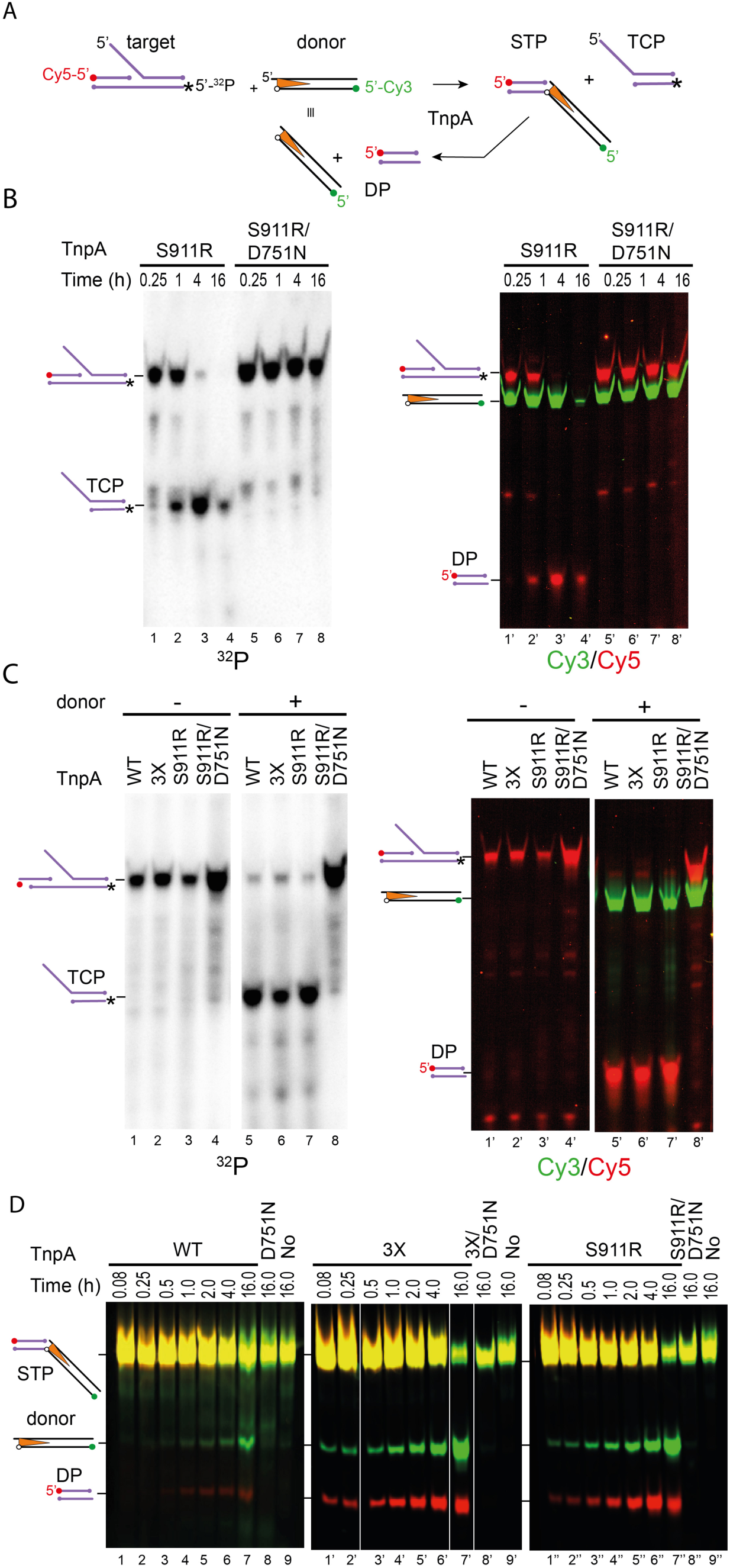
Turnover of the integation reaction *in vitro*. (A) Integration of a pre-cleaved 5’ Cy3-labeled donor IR (green dot) into a DNA fork that is 5’ Cy5-labeled on the leading strand (red dot) and 5’ ^32^P- labeled on the leading strand template (*) generates a radiolabeled target cleavage product (TCP) and a fluorescently labeled strand transfer product (STP) which is in turn cleaved back to regenerate the initial donor end and release a secondary disintegration product (DP) corresponding the left moiety of the initial fork. (B) Time course analysis performed with TnpA^S911R^ and TnpA^S911R/D751N^ showing the disappearance of the initial target substrate and accumulation of the TCP and DP over time. (C) Prolonged reaction (16h) performed with the indicated TnpA proteins in the presence (+) or absence (-) of the donor IR. No TCP or DP accumulates in the absence of donor demonstrating that integration is a pre-requisite for disintegration. (D) Kinetics of disintegration. A doubly Cy5- and Cy3-labeled STP was incubated with TnpA^WT^, TnpA^3X^, TnpA^S911R^ and the corresponding D751N catalytic mutants for the indicated timepoints to follow donor and DP accumulation over time.

These results show that DNA fork targeting by TnpA is an efficient process that can lead to complete depletion of the target after multiple rounds of integration-disintegration *in vitro*. They also suggest that additional factors are required *in vivo* to efficiently covert strand transfer intermediates into final replicative transposition products.

## DISCUSSION

### A replication hijacking mechanism for replicative transposition

The movement of all transposable elements requires that reaction intermediates be processed by the host to be effective (1,5). In particular, specific transposons exploit host DNA replication to produce new copies of themselves each time they move, which allows them to multiply faster than their host and thus to optimize their spread and dispersal. A classical mechanism of replicative transposition is termed “paste-and-copy” or “copy-in” because replication of the transposon is concomitant with its integration into the target (Fig. 7). This mechanism has been studied in detail for bacteriophage Mu and was extrapolated to other elements which like Mu and Tn3-family transposons, generate cointegrate transposition products (11,42,43).

**Figure 7.**
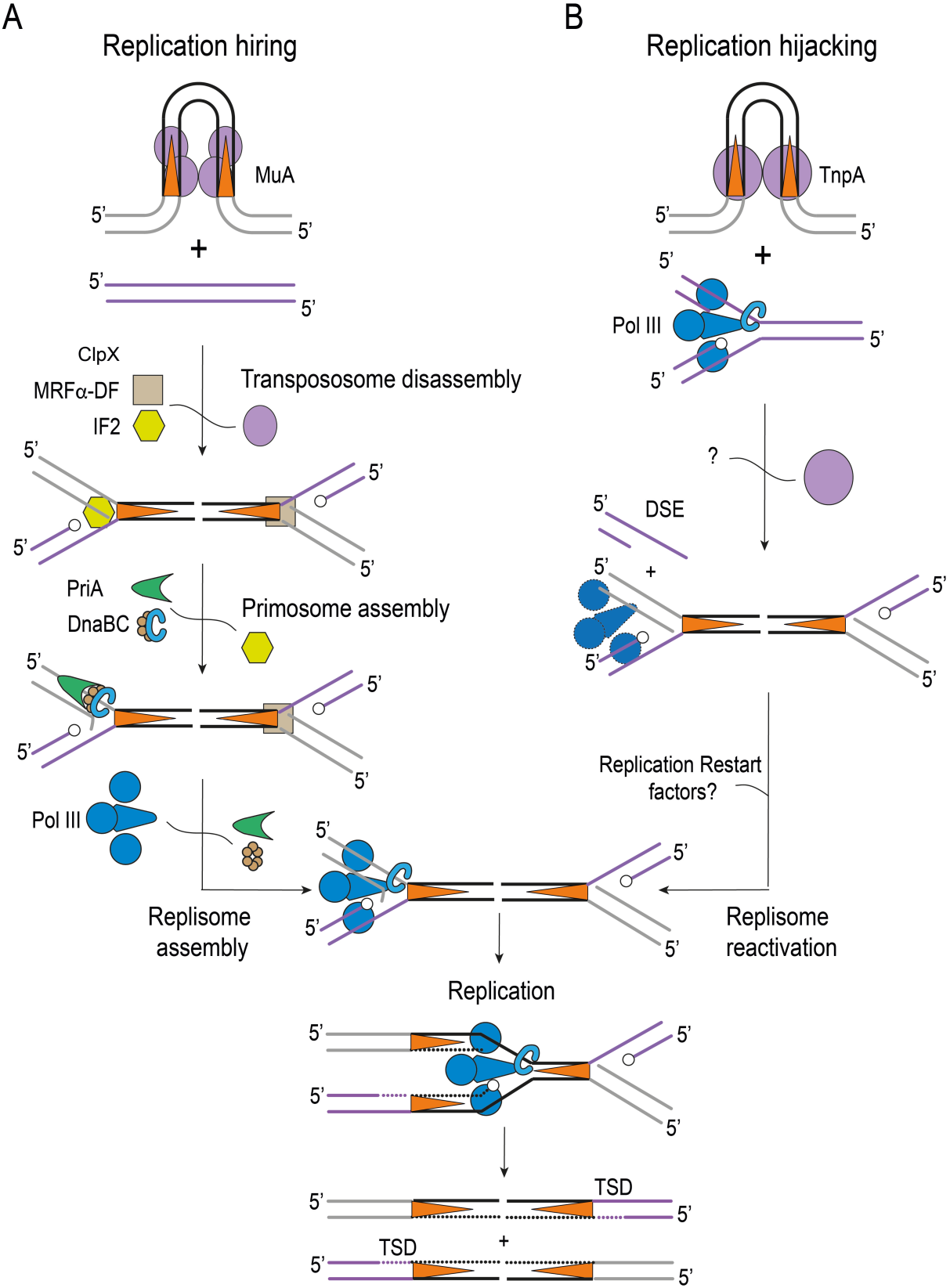
Alternative models for replication recruitment during copy-in replicative transposition. (A) Classical replication hiring mechanism exemplified by bacteriophage Mu. A tetramer of the MuA transposase (purple oval) helped by the target selector MuB (not shown) concertedly joins both Mu ends (orange triangles) to 5-bp staggered positions of a target DNA duplex generating a double-fork strand transfer intermediate. The Mu transpososome is then disassembled in an ATP-dependent reaction involving successive host factors: the ClpX chaperone, a yet unidentified Mu replication- disassembly factor MRFα-DF and the translation initiation factor IF2. IF2 Binding at one end of Mu recruits the PriA/PriC/DnaT restart system, which in turn gives access to the helicase loading complex DnaB/DnaC. Loading of the replicative DnaB helicase onto the lagging strand template promotes assembly of DNA polymerase III holoenzyme and replication of Mu genome by the replisome. (B) Replication hijacking model proposed here for Tn*4430* and Tn*3*-family transposons. A TnpA dimer directly transfers the transposon ends at the level of an ongoing replication fork within the target. Transfer of both ends downstream of leading strand 3’OH end generates a double-strand end (DSE) that is either degraded or repaired by homologous recombination. Replisome reactivation leads to transposon duplication. The transposon DNA is shown in black, the donor in grey and the target in purple. Open circles indicate the position of free 3’OH ends. Newly synthesized DNA strands are presented by dashed lines. TSD, target site duplication

Replicative transposition of Mu follows a “replication hiring” mechanism during which the cell replication apparatus is recruited at the level of integration intermediates after strand transfer catalyzed by the transposase MuA (Fig. 7A). Replication initiation of Mu is an ATP consuming multistep process requiring the action of the chaperone ClpX and a set of host factors (*e.g*., MRFα−DF and IF2) to first displace the transposition complex from the strand transfer product, and then promote PriA-dependent assembly of the replisome at one end of the Mu genome (Fig. 7A) (22,23,44). This process is thus reminiscent of ectopic replication initiation which takes place during DNA repair and replication re-start at stalled replication forks (22,24,33,45,46).

By contrast, our data suggest that Tn*3*-family transposons use an alternative “replication hijacking” mechanism by recruiting the replisome directly from ongoing replication within the target molecule (Fig. 7B). An immediate consequence of this mechanism is the finding that target replication is a requirement for Tn*4430* transposition, as was initially shown for Tn*1* using replication-defective λ phage derivatives (47). Preventing replication of the donor molecule has no impact on transposition, whereas blocking the initiation of replication in the target abolishes it completely. In contrast, neither donor nor target replication is required for transposition during lytic growth of Mu, which is consistent with the fact that replication is recruited after strand transfer in this case (45,48,49). However, collision between the Mu transposition complex and the replisome has been proposed to repair strand transfer intermediates resulting from primary integration of Mu into the host genome to establish lysogeny, albeit this is a non-replicative process (50).

Replisome recruitment by recurrent “hiring” mechanism is crucial for Mu lytic growth since it undertakes up to ∼100 rounds of replication, while the infected cells themselves do not replicate (11,21,51). In contrast, conventional transposons, which typically move at a low frequency, must ensure that transposition only occurs occasionally and under optimal conditions. In our “replication hijacking” model, transposition directly depends on ongoing replication providing a means to sense the physiological state of the host cells and favor transposition in bacterial populations that undergo expansion and/or frequent movements of other mobile genetic elements such as conjugative plasmids. This would increase the chances of efficient transmission, both vertically and horizontally, and should be taken into account in strategies to combat the spread of antibiotic resistance. Direct use of the replication apparatus also synchronizes transposition catalysis with replication ensuring efficient transposition while avoiding the formation of partially processed and potentially harmful intermediates. It remains to be determined whether other elements that transpose with the “copy-in” mode recruit replication via the Mu “replication hiring” pathway or through “replication hijacking” as proposed here for the Tn3 family (51).

### Interaction between the transposition complex and replication?

Alignment of integration sites revealed that Tn*4430* inserts with little sequence specificity, apparently recognizing a weak consensus sequence 5’-[G/C][T/A]nnn[A/T][G/C]-3’ overlapping the 5-bp TSD. However, this consensus is not the only determinant of Tn*4430* insertion specificity. In particular, the presence of several copies of the *lacO* operator in a target plasmid altered the regional preference of the transposon irrespective of the target sequence, creating a strong hotspot between the unidirectional replication origin of the target and the *lacO* array, and changing the direction of replication of the target plasmid further altered the integration pattern. We propose that the pseudo-palindromic *lacO* sequences would somehow interfere with replication fork progression (*e.g*., by forming DNA secondary structures), thereby promoting the accumulation of preferred target DNA structures in the hotspot region. Pausing of replication could allow TnpA to catalyze strand transfer into the target before replication processing when the replisome restarts (Fig. 7B). Consistent with direct targeting of DNA replication, several Tn*3*-family members were shown to preferentially insert into or near the replication origin of target plasmids (52,53); whereas chromosomal integration of Tn*917* showed a strong bias for the terminus region where replication forks meet, collapse and eventually reassemble to complete replication (33,54,55).

A model for target DNA binding by Tn4430 TnpA was obtained from the cryo-EM structure of a post-integration complex showing a dimer of hyperactive TnpA^S911R^ bound onto paired strand transfer products mimicking the transfer of the 3’ ends of Tn4430 into the target DNA (15). Although the target branch of the substrate was poorly structured in this complex, modeling the density map suggests that the target DNA occupies a groove that crosses the TnpA dimer from one side to the other. In this model, the DNA is sharply bent to fit the groove and position the strand transfer sites 5 bp apart in the active TnpA dimer (15). Our results show that Tn*4430* TnpA has little or no affinity for conventional DNA duplexes, but that it binds to specific DNA structures mimicking replication intermediates and can use them as a substrate for strand transfer *in vitro*. The preferred target appears to be a Y-shaped DNA molecule with the structure of a simplified replication fork. Strand transfer occurred at a specific position, joining the transposon end to the template strand, immediately downstream of the 3’OH group of the leading strand. *In vivo*, this configuration would lead to duplication of the transposon as DNA synthesis resumes or continues (Fig. 7B). However, the transfer of only one end has been observed under conditions that were previously shown to hyper- activate TnpA (*i.e*., in the presence of Mn^2+^ instead of Mg^2+^)(13). Since Tn*4430* transposition generates 5-bp TSDs, joining of the second end is expected to take place 5 base pairs upstream, in the template of the lagging strand (Fig. 7B). In addition, the strand transfer products proved to be unstable being cleaved back by TnpA, leading to abortive re-excision (*i.e*, disintegration) of the transposon end. This suggests that the target configuration used *in vitro* is likely not optimal for catalysis of concerted end joining and that an additional actor is required to carry out the reaction and drive it forward *in vivo*. This actor could be a host factor or DNA synthesis itself, which would further reinforce the functional coupling between transposition and replication. Both processes might also overlap, with one transposon end being transferred before the other and replication restart taking place between the two as was proposed in early asymmetric models for replicative transposition (56,57). Recent studies have shown that replisomes are dynamic multi-protein machines with far greater flexibility than previously appreciated (58–61). Regardless of the replication stage targeted by by Tn*4430*, this flexibility is likely to allow the transposition complex to hijack a replication fork and come into contact with the appropriate DNA structure for integration.

Interestingly, a functional link between transposition and DNA replication has been reported for an increasing number of transposable elements in prokaryotes and eukaryotes, but the exact significance of this interaction often remains unclear (62).

Several transposons appear to preferentially integrate into the lagging strand template which is thought to provide a window of “vulnerability” to facilitate access to DNA (63). For the “cut-and-paste” transposon Tn*7*, which normally does not require DNA replication, targeting of lagging strand synthesis is proposed to favor transposition into mobile genetic elements such as conjugative plasmids, thereby enhancing the horizontal dispersal of the element (64). For IS*608* and other members of the IS*200*/IS*605* family, lagging strand synthesis generates stretches of ssDNA that are the substrates for both the excision and integration reactions catalyzed by the HUH transposase (65,66). Specific Group II mobile introns integrate downstream of Okasaki fragments to use the free 3’OH end to prime reverse transcription of the intron RNA (9). Interestingly, genome-scale analyses suggest that this strategy has been inherited by L1, the most prevalent transposable element in human (67–69).

For some of these elements, directing the transposition complex to replicating targets appears to depend on interactions with replication-specific proteins such as the β-sliding clamp or PCNA processivity factors (66,70–75). Intriguingly, the transposase of the Tn*3*-family transposon Tn*1721* was recently reported to interact with the β-clamp through a specific motif buried in the N-terminal DNA binding domain of the protein (75). Since this motif is only conserved in a subset of Tn*3*-family members, including Tn*4430*, more work is needed to fully assess the significance of β-clamp and possibly other replication partners in Tn*3*-family targeting mechanism.

## Acknowledgement

We are grateful to P. soumillion, P. Hols, C. Claeys Bouuaert and all members of the BGM group at the LIBST institute of UCLouvain for fruitful discussions and advices and for crital reading of the manuscript.

## Data Availability Statement

Additional raw data underlying this article including mapping of Mini-TnKm insertion sites into pGB2Ts derivatives, transposition frequencies measurements, EMSA quantifications and uncropped Electrophoresis gels are available at https://zenodo.org/records/20465000. All other relevant data are in the article or the Supplementary data file.

## Supplementary figures and tables

**Figure S1.**
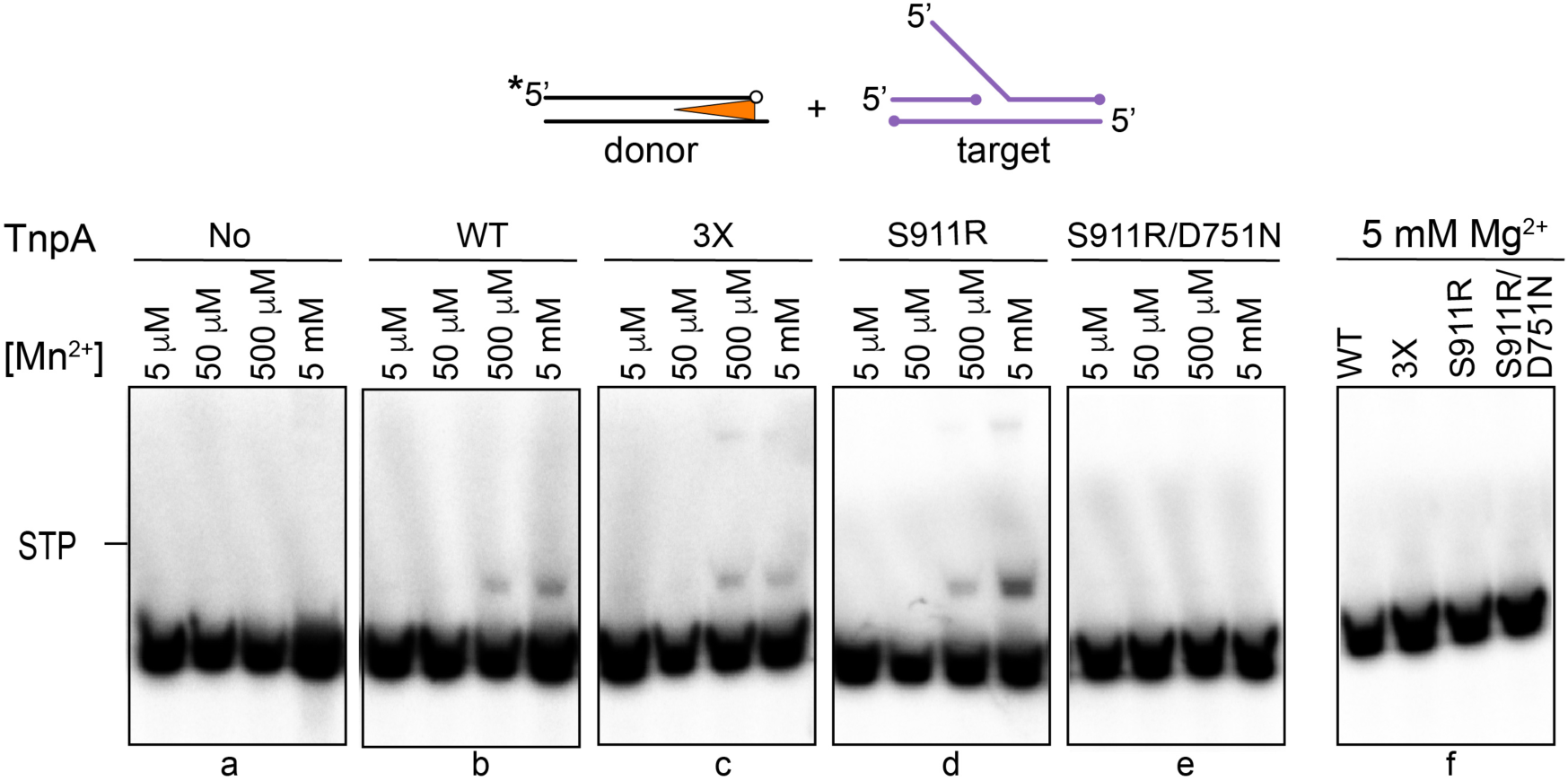
Cofactor requirement for integration *in vitro*. Integration of a 5’ P^32^-labeled pre-cleaved IR end into an fork-like target was performed with the indicated variants of TnpA in the presence of increasing concentrations of Mn^2+^ (from 5 μM to 5 mM, panels a-e) or 5 mM of Mg^2+^ (panel f). Strand transfer products (STP) were visualized by phosphorimaging following electrophoresis in a 7.5 % polyacrylamide gel. No, no TnpA.

**Figure S2.**
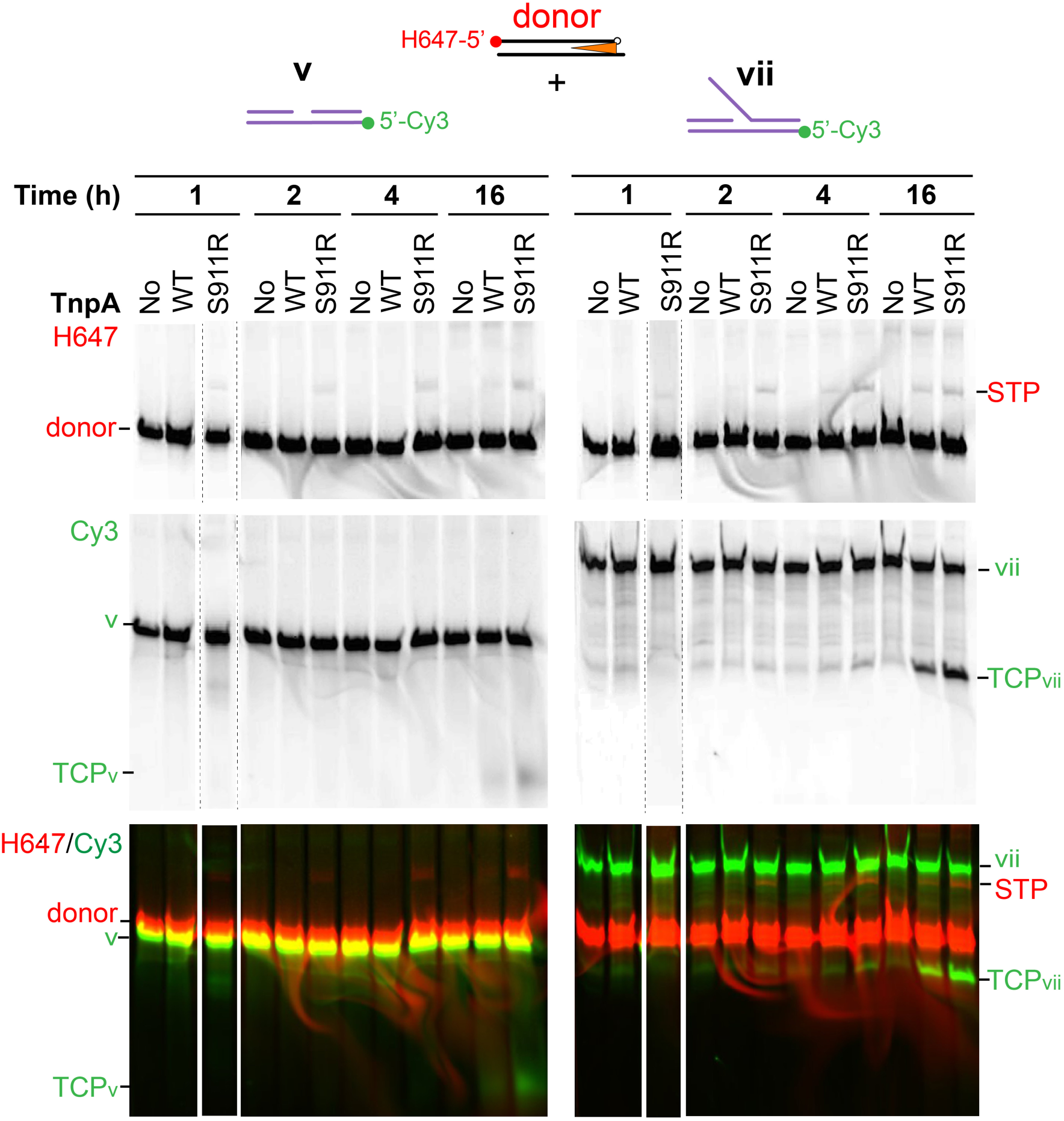
Differential kinetics of STP and TCP accumulation. Integration reactions of a 5’ H647- labeled pre-cleaved donor IR into the Cy3-labeled gapped linear substrate v (left) and the fork-like substrate vii (right) were performed in the presence of TnpA^WT^ or TnpA^S911R^ for the indicated time and analyzed by electrophoresis in 7.5 % polyacrylamide gels. The gels were scanned for H647 fluorescence to detect strand transfer products (STP, upper panels) and for Cy3 fluorescence to detect target cleavage products (TCP, middle panels). The bottom panel shows the overlay of both signals. No, No TnpA.

**Figure S3.**
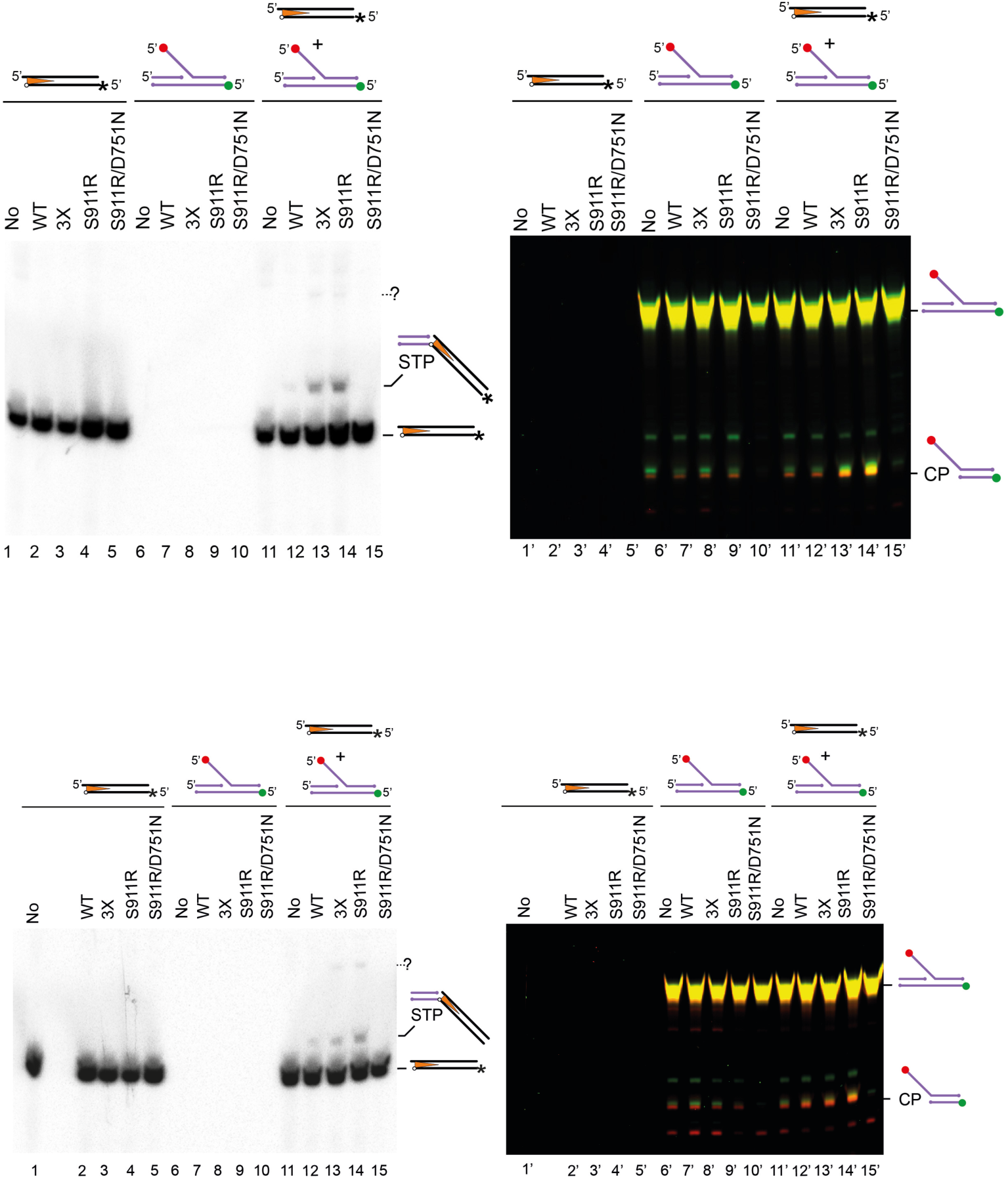
Integration into a DNA fork requires both the donor and target partners. Integration of a pre-cleaved 5’ ^32^P-labeled (*) IR end into a Cy5- (red dot) and Cy3- (green dot) labeled DNA fork was carried out with the indicated TnpA proteins as described in Figure 5. As a control, reactions were performed with the donor or the target alone as depicted above the electrophoretic gel. The gel was analyzed by phosphorimaging (left) and fluorescence scanning (right) to detect radiolabeled strand transfer products (STP) and target cleavage products (TCP), respectively. Specific products were only observed in reactions where both partners were incubated together with the active transposases. “?” refers to a minor product observed with TnpA^3X^ and TnpA^S911R^ that could not be characterized further. Upper part of the figure corresponds to the experiment of Figure 5 and lower part comes from an independent experiment.

**Figure S4.**
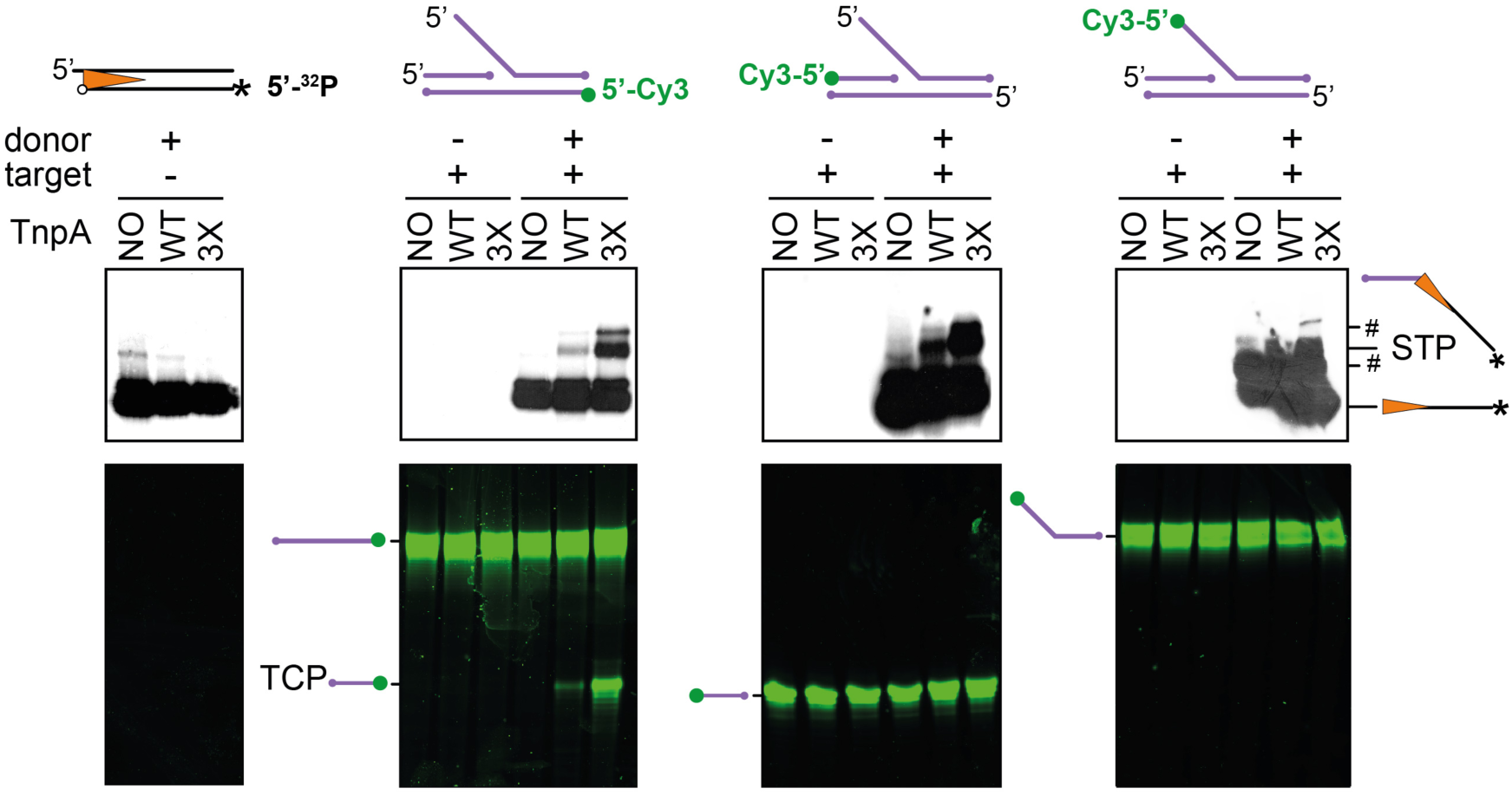
Target strand specificity of strand transfer. TnpA^WT^ and TnpA^3X^-mediated Integration of a pre-cleaved 5’ ^32^P-labeled (*) IR end into forks that have been separately Cy3-labelled at the 5’ end of each DNA strand (green dot). Reactions were run on an 8% denaturing gel and analyzed by phosphorimaging to identify ^32^P-labeled strand transfer products (STP), and by Cy3 fluorescence scanning to determine which DNA strand is cleaved to form target cleavage products (TCP). Reactions were performed in the absence of the target or donor substrate to rule out possible products arising from simple processing of the IR end or the DNA fork, respectively. Hashtag symbol (#) indicates the position of radioactive bands corresponding to incompletely denatured donor or STP, respectively.

**Figure S5.**
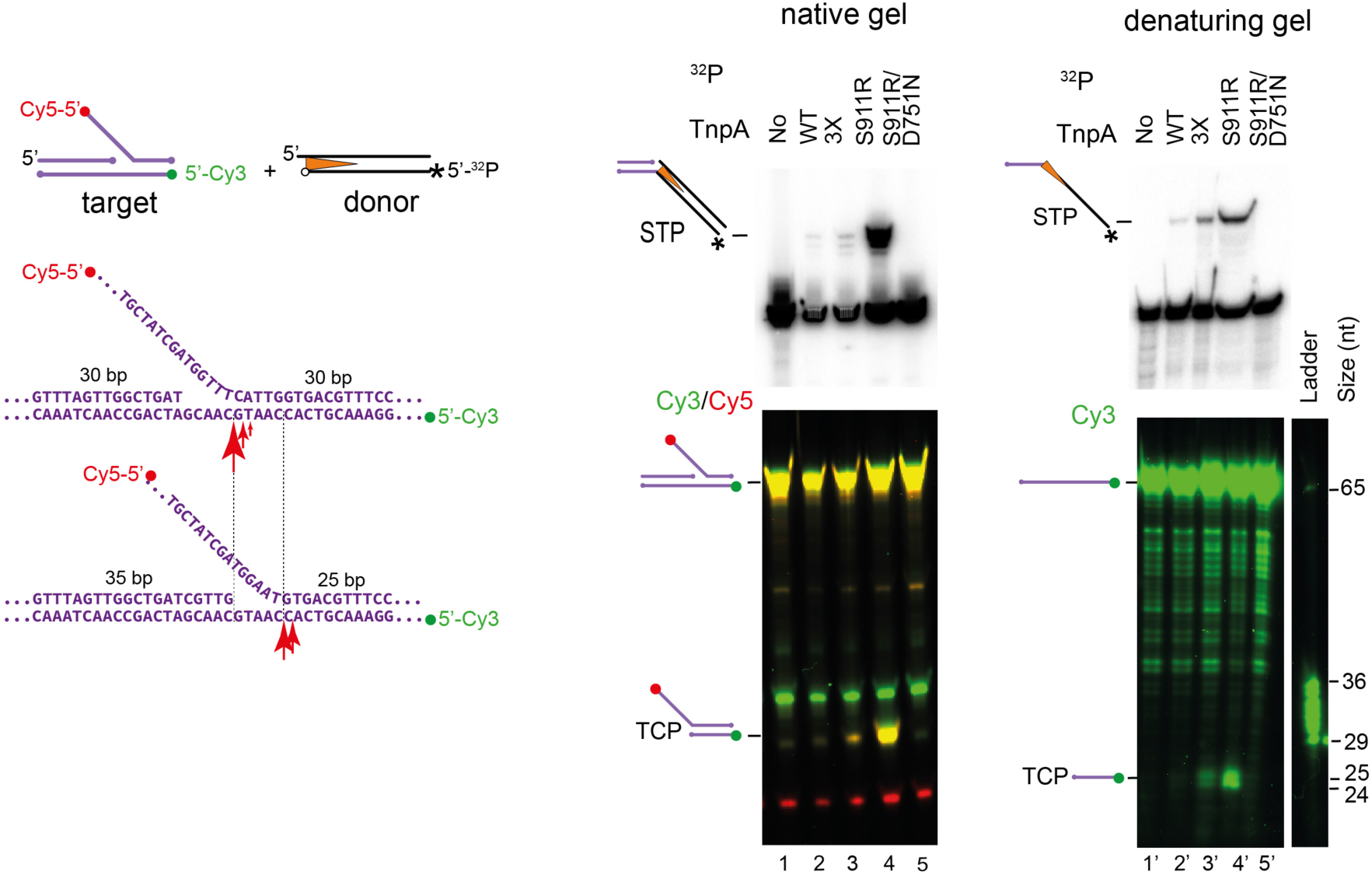
Structural determinant of integration. Integration of a P^32^-labelled donor IR was examined into an asymmetric Cy5/Cy3-labeled DNA fork in which the branchpoint is shifted 5 bp compared to fork analyzed in Figure 5 (vertical dashed lines). Mapping of the the cleavage sites reveals that strand transfer occurred at, or adjacent to the branch point as was observed for the symmetrical fork (red arrowheads).

**Table S1.**
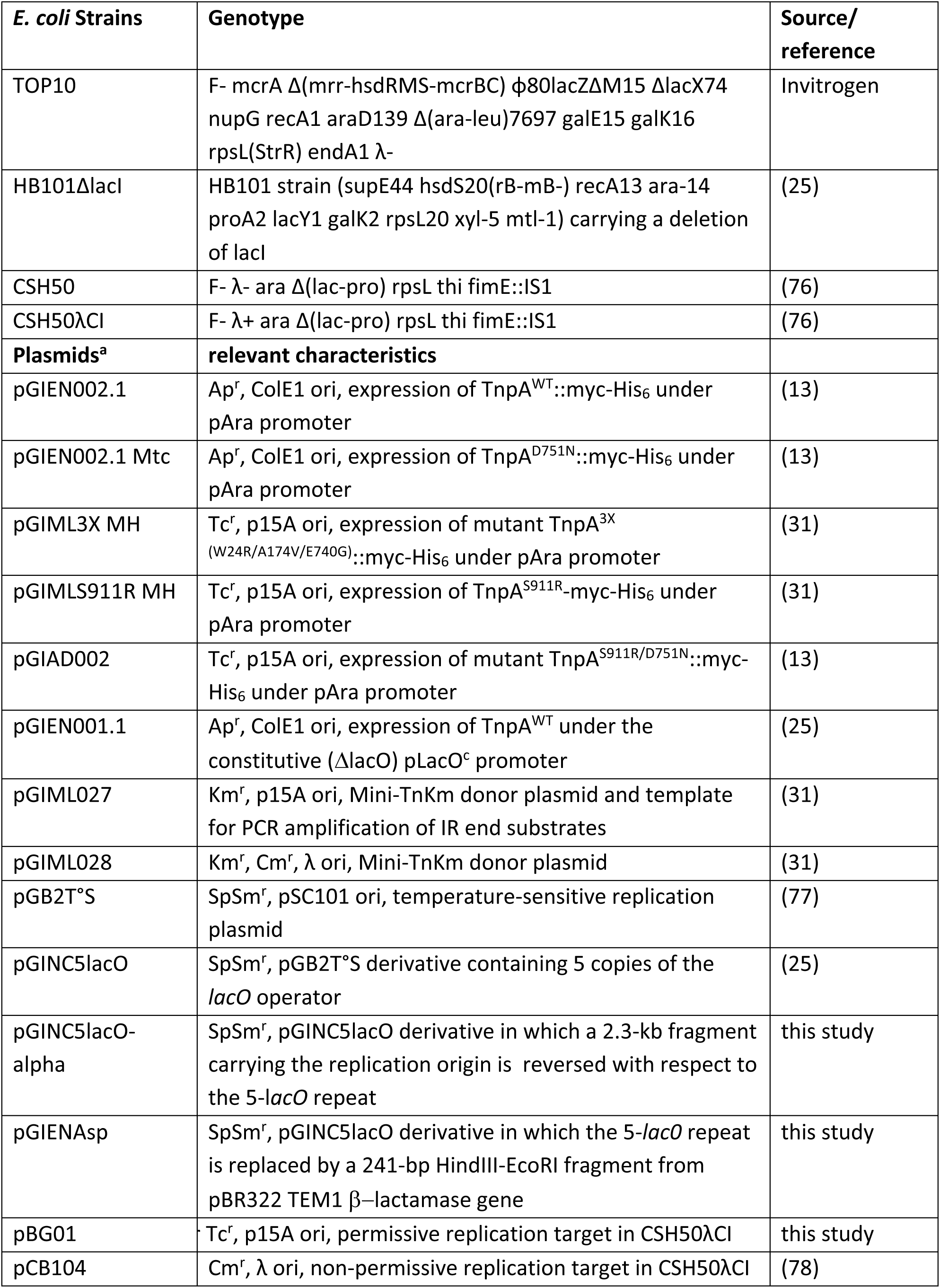
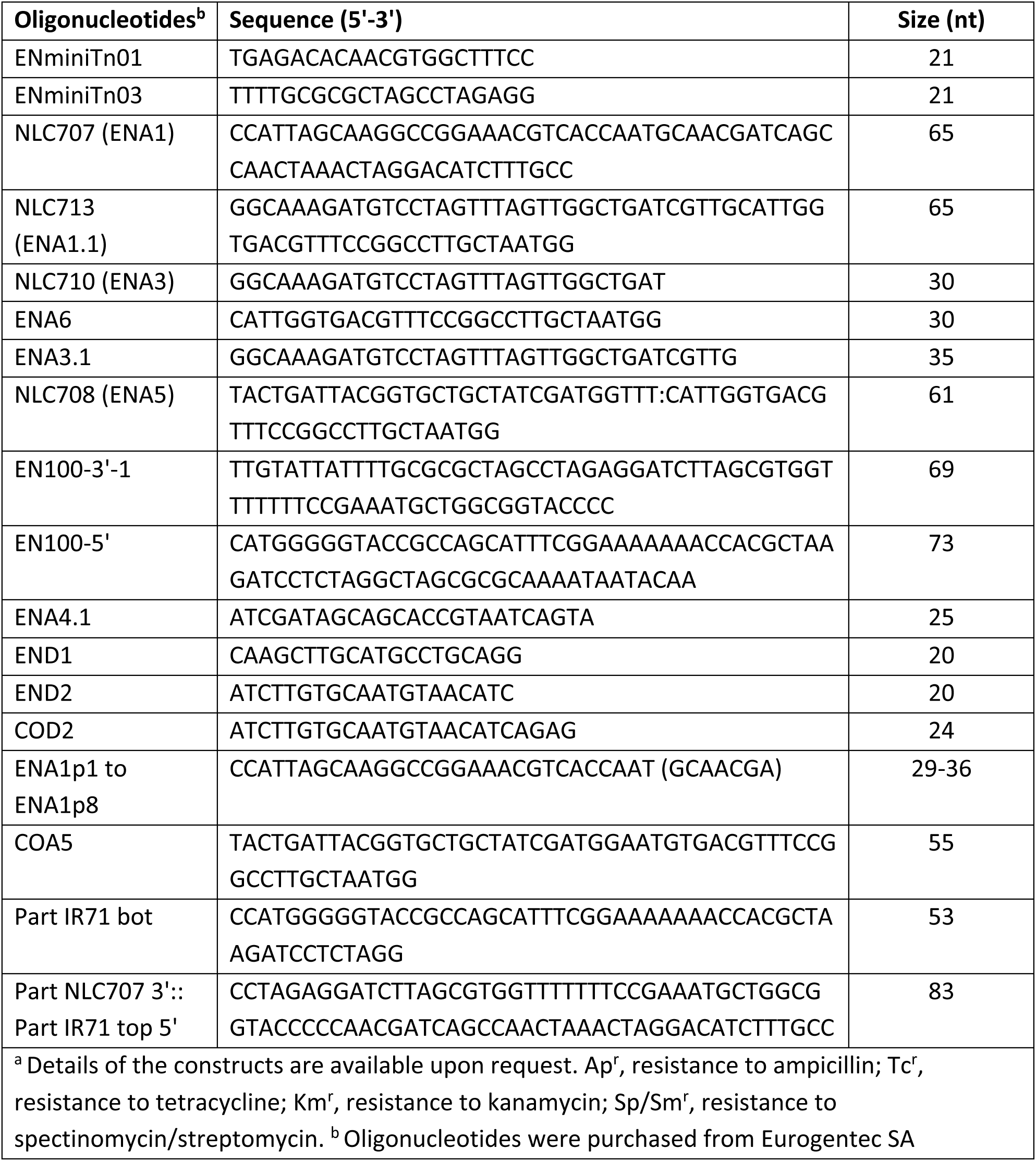
Bacterial strains, plasmids, and oligonucleotides used in this study.

**Table S2.** DNA substrates.

| Figure | Substrate | Oligonucleotides assembly - PCR |
| --- | --- | --- |
| 3, 4, S2 | Target |  |
|  | i | 5'**Cy3**-NLC707 |
|  | ii | 5'**Cy3**-NLC707, NLC713 annealed |
|  | ii | 5'**Cy3**-NLC707, NLC710 annealed |
|  | iv | 5'**Cy3**-NLC707, ENA6 annealed |
|  | v | 5'**Cy3**-NLC707, NLC710 annealed |
|  | vi | 5'**Cy3**-NLC707, ENA3.1 annealed |
|  | vii | 5'**Cy3**-NLC707, NLC710, NLC708 annealed |
|  | vii | 5'**Cy3**-NLC707, NLC708, ENA 4,1 annealed |
| 4, S2 | IR donor | 5'**H647**-EN100-3'-1, EN100-5' annealed |
| 5, S1, S3, S4, S5 | IR donor | PCR with 5'**P<sup>32</sup>**-COD2 and END1 on pGIML027 + NcoI digestion |
| 5, S1, S3 | Fork | 5'**Cy3**-NLC707, NLC710, 5'**Cy5**-NLC708 annealed |
| S4 | Fork | 5'**Cy3**-NLC707, NLC710, NLC708 annealed<br>NL707, 5'**Cy3**-NLC710, NLC708 annealed<br>NL707, NLC710, 5'**Cy3**-NLC708 annealed |
| S5 | Fork | 5'**Cy3**-NLC707, ENA 3.1, 5'**Cy5**-COA5 annealed |
| 5, S5 | ladder | 5'**Cy3**-ENA1p1 to -ENA1p8 |
| 6 | IR donor | PCR with 5'**Cy3**-END2 and END1 on pGIML027 + NcoI digestion |
|  | Fork | 5'**P32**-LC707, NLC710, 5'**Cy5**-NLC708 annealed |
|  | STP | 5'**Cy5**-NLC710, Part IR71 bot,<br>5'**Cy3**-Part NLC707 3':: Part IR71 top 5' annealed |
| 5'-Hylite 647 (**H647**), 5'-Cyanidine 5 (**Cy5**) or 5-Cyanidine 3 (**Cy3**) -labelled oligonucleotide were purchased from Eurogentec SA. 5'-P<sup>32</sup> labelling was performed with polynucleotidyl kinase and (γ32P)ATP. |  |  |

